# Hofbauer cells and fetal brain microglia share transcriptional profiles and responses to maternal diet-induced obesity

**DOI:** 10.1101/2023.12.16.571680

**Authors:** Rebecca Batorsky, Alexis M. Ceasrine, Lydia L. Shook, Sezen Kislal, Evan A. Bordt, Benjamin A. Devlin, Roy H. Perlis, Donna K. Slonim, Staci D. Bilbo, Andrea G. Edlow

**Author notes:** These authors contributed equally (Rebecca Batorsky, Alexis M. Ceasrine, Andrea G. Edlow). Correspondence: Andrea G. Edlow, Massachusetts General Hospital, Vincent Center for Reproductive Biology 55 Fruit Street, Thier Research Building, 9^th^ floor Boston, MA 02114.

## Abstract

Maternal immune activation is associated with adverse offspring neurodevelopmental outcomes, many mediated by in utero microglial programming. As microglia remain inaccessible throughout development, identification of noninvasive biomarkers reflecting fetal brain microglial programming could permit screening and intervention. We used lineage tracing to demonstrate the shared ontogeny between fetal brain macrophages (microglia) and fetal placental macrophages (Hofbauer cells) in a mouse model of maternal diet-induced obesity, and single-cell RNA-seq to demonstrate shared transcriptional programs. Comparison with human datasets demonstrated conservation of placental resident macrophage signatures between mice and humans. Single-cell RNA-seq identified common alterations in fetal microglial and Hofbauer cell gene expression induced by maternal obesity, as well as sex differences in these alterations. We propose that Hofbauer cells, which are easily accessible at birth, provide novel insights into fetal brain microglial programs, and may facilitate the early identification of offspring vulnerable to neurodevelopmental disorders in the setting of maternal exposures.

## Introduction

Microglia play a key role in neurodevelopment by modulating synaptic pruning, neurogenesis, phagocytosis of apoptotic cells, and synaptic plasticity^1–4^. Aberrant programming of fetal microglia in the setting of maternal immune activation has accordingly been identified as a key mechanism underlying abnormal fetal brain development^5–9^ and likely contributes to the pathogenesis of neurodevelopmental and psychiatric disorders^10–13^. In part because microglia remain inaccessible in fetal life and postnatally, there is currently no way to identify which offspring may be most at risk for adverse neurodevelopmental and psychiatric morbidity after *in utero* exposures. Methods of determining whether or how *in utero* exposures may have primed fetal microglia could facilitate intervention during critical developmental windows when outcomes can potentially be modified.

Precursors of many tissue-resident macrophages, including microglia, originate in the fetal yolk sac^14–16^. As the yolk-sac is a pre-placental structure, yolk-sac-derived macrophages likely also colonize the placenta, where they are called Hofbauer cells, but prior lineage-tracing has not explicitly focused on the placenta. These cells share exposure to the same intrauterine environment as microglia, and their connection to the developing brain has been posited as a result of their role in the maternal-to-fetal transmission of neurotropic viruses such as Zika, cytomegalovirus, and HIV^17–21^. In support of commonality between these cell types, prior work by our groups has demonstrated that maternal immune activation in the setting of high-fat diet primes both placental macrophages and fetal brain microglia toward a highly-correlated, pro-inflammatory phenotype^22,23^. Still, whether Hofbauer cells manifest the same transcriptional programs as fetal microglia has not yet been investigated.

To address these gaps in understanding, we sought to definitively trace Hofbauer cells from the fetal yolk sac to the placenta. Using an inducible macrophage reporter mouse model, we demonstrate that yolk sac-derived macrophages comprise the majority of tissue resident macrophages in both placenta and brain in late embryonic development (embryonic day 17.5; e17.5). Further, we isolated placental resident macrophages and microglia from fetuses of diet-induced obese and control dams (wild-type C57BL/6J) at e17.5 and characterized cells using single cell RNA-Sequencing (scRNAseq). Using X– and Y-chromosome markers, we distinguished fetal placental macrophages, or Hofbauer cells, from placenta-associated maternal monocyte/macrophage (PAMM) populations, generating new insights into transcriptional differences between Hofbauer cells and PAMMs both at baseline and in the setting of maternal diet-induced obesity. Specifically, we identified subpopulations of fetal placental macrophages that transcriptionally mirror fetal brain macrophages and found that maternal obesity affected gene expression similarly in both microglia and Hofbauer cells, both in terms of differentially expressed genes and biological processes and pathways. Functional analyses of differentially expressed genes provided new insights into sex differences in immune function and the response to maternal obesity in both placental and fetal brain macrophages.

Taken together, these data suggest that late pregnancy placental macrophages reflect the programs of fetal brain microglia, both at baseline and in response to the perturbation of maternal diet-induced obesity. Hofbauer cells therefore have the potential to provide novel insights into the relationship between placental and brain immune programming in the setting of specific *in utero* exposures such as maternal diet-induced obesity. These more accessible fetal placental macrophages may help identify offspring most vulnerable to neurodevelopmental morbidity in the context of maternal exposures such as obesity.

## Results

### Tissue-resident placental macrophages are yolk sac derived

In both mice and humans, the maternal-fetal interface is comprised of the maternally-derived decidua and the fetally-derived placenta. Fetal placental macrophages have been identified as early as 18 days post conception in humans, and 10 days post conception in mice (e10) prior to the full vascularization of the placenta^24^. This suggests that extra-embryonic placental macrophages (Hofbauer cells) likely derive from the yolk sac, as do many other tissue-resident macrophages, including microglia^25^.

To determine whether fetal placental macrophages are in fact yolk sac-derived, we crossed transgenic mice carrying a floxed Rosa-*tdTomato* allele^26^ with tamoxifen inducible transgenic *Csf1R-Cre^ER^*mice to permanently label yolk sac progenitors. Csf1r is active in yolk sac progenitor cells at gestational day 8-9 (gd8-9), thus a 4-hydroxytamoxifen (4-OHT) pulse at gd8.5 will label yolk sac-derived macrophages prior to their migration out of the yolk sac^27^. Migration of macrophages out of the yolk sac to colonize the fetal brain and other tissues begins around e9^14^, and Csf1r+ placental macrophages are detectable beginning at e10^24^. Thus, we delivered 4-OHT to Cre^ER^ negative dams (tdTomf/+ or tdTomf/f) crossed with Cre^ER^ positive sires, so that 4-OHT only induced *Csf1R-Cre^ER^* activity in *fetal* macrophages within the embryos (Figure 1A). We then assessed tdTomato+ cells at e17.5 in placenta and brain from Cre^ER^-negative and Cre^ER^-positive embryos in conjunction with immunohistochemistry for ionized calcium binding adaptor molecule (Iba1), a marker for macrophages. In the placenta labyrinth, we saw robust colocalization of tdTomato and Iba1 at e17.5 (approximately 83% of Iba1+ cells were also tdTomato+; Figure 1B, C’). As expected, we also detected substantial colocalization in e17.5 brain (hippocampus shown in Figure 1B, D). We did not see any tdTomato signal in Cre negative placenta or hippocampus (Figure 1 C, D). Together, these data suggest that resident placental macrophages are primarily derived from yolk sac progenitor cells, similar to fetal microglia. Additional staining demonstrated that placental macrophages were found within the villous tissue and were not primarily located within the vasculature (Figure S1).

**Figure 1.**
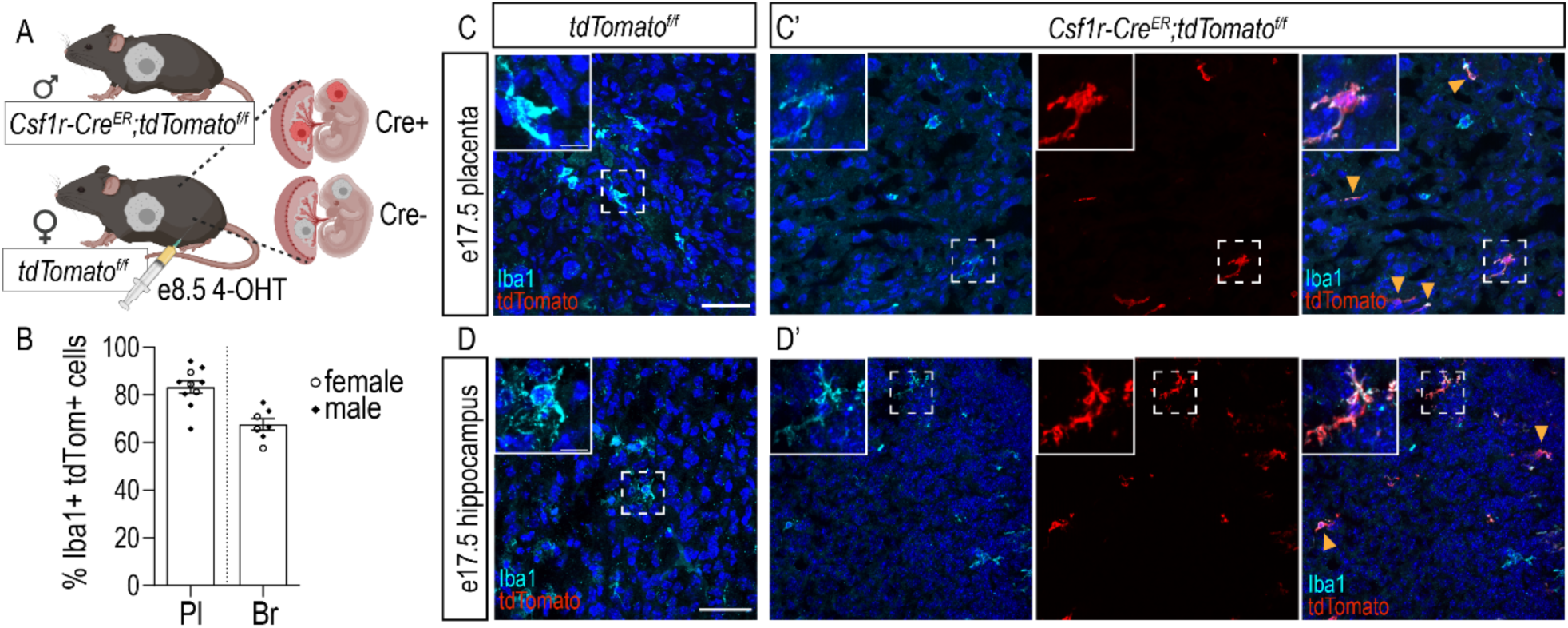
Placental macrophages are yolk sac derived. **A.** Schematic of fetal yolk sac macrophage labeling. Briefly, male *Csf1R-Cre^ER^;tdTomato^f/f^* mice were timed-mated to female *tdTomato^f/f^* mice. Pregnant females were injected with 4-hydroxytamoxifen (4-OHT) at gestational day 8.5. Embryos were collected at embryonic day 17.5. **B.** Percent of macrophages (Iba1+ cells) labeled with tdTomato in embryonic placenta and hippocampus following 4-OHT administration at e8.5. Open circles represent individual female embryos and closed diamonds represent individual male embryos (n=4 litters). Pl = placenta; Br = brain (hippocampus) **C-C’.** Representative images of Iba1 and tdTomato in control (C) and reporter (C’) placenta from e17.5 embryos. **D-D’.** Representative images of Iba1 and tdTomato in control (D) and reporter (D’) hippocampus from e17.5 embryos. Arrowheads indicate double-positive (Iba1+ tdTomato+) macrophages/microglia in reporter tissue. Scale 50µm, inset scale 10µm.

### Fetal placental and brain macrophages are heterogeneous populations with shared cluster-specific signatures

To identify subpopulations of fetal placental and brain macrophages, we performed single-cell RNA-sequencing (10X Genomics) on macrophage-enriched single-cell suspensions from matched placenta and fetal forebrain tissue from both male and female embryos at e17.5 (Figure S2A). Matched placenta and fetal brains were collected from 17 mouse embryos, comprising 8 embryos from obese dams (4 male, 4 female) and 9 embryos from control dams (4 male, 5 female). 197,000 cells were sequenced to an average depth of approximately 22,000 reads/cell. In order to identify cell types present in our data, we used graph-based clustering followed by identification of cluster-specific marker genes and comparison of marker gene expression and average-expression profiles of all clusters with published placental^28–32^ and fetal brain^33–35^ datasets (see Methods, Figure S2B-C, Supplemental Table 1). We selected macrophage and monocyte-like clusters for our main analysis in light of recent evidence of monocyte-to-macrophage transitional populations at the maternal-fetal interface^36^. The final clusters are visualized as uniform manifold approximation and projection (UMAP) plots for brain (Figure 2A) and placenta (Figure 2B) along with expression plots demonstrating the top three marker genes per cluster. Clusters have been named with a cell type prefix (Mg: microglia; HBC: Hofbauer cell; Mono_FBr: fetal brain monocytes; Mono_FPl: fetal placental monocytes; PAMM: placenta-associated maternal macrophages and monocytes) followed by the top marker gene in the cluster. Clusters primarily engaged in cell cycle functions (e.g., processes integral to DNA replication) end in _cellcycle. A complete list of cluster marker genes can be found in Supplemental Table 1. Interrogation of the transcriptional signatures of brain clusters revealed two distinct groupings of microglia. One grouping was defined by a more robust yolk sac signature, defined based on the signature of yolk-sac derived macrophages^37,38^ (Figure 2C). We designated these clusters “yolk sac imprint Microglia”, Mg_YSI, which are further divided into two subpopulations, MgYSI_Pf4 and MgYSI_cell cycle; the latter represents the cell cycling component.

**Figure 2.**
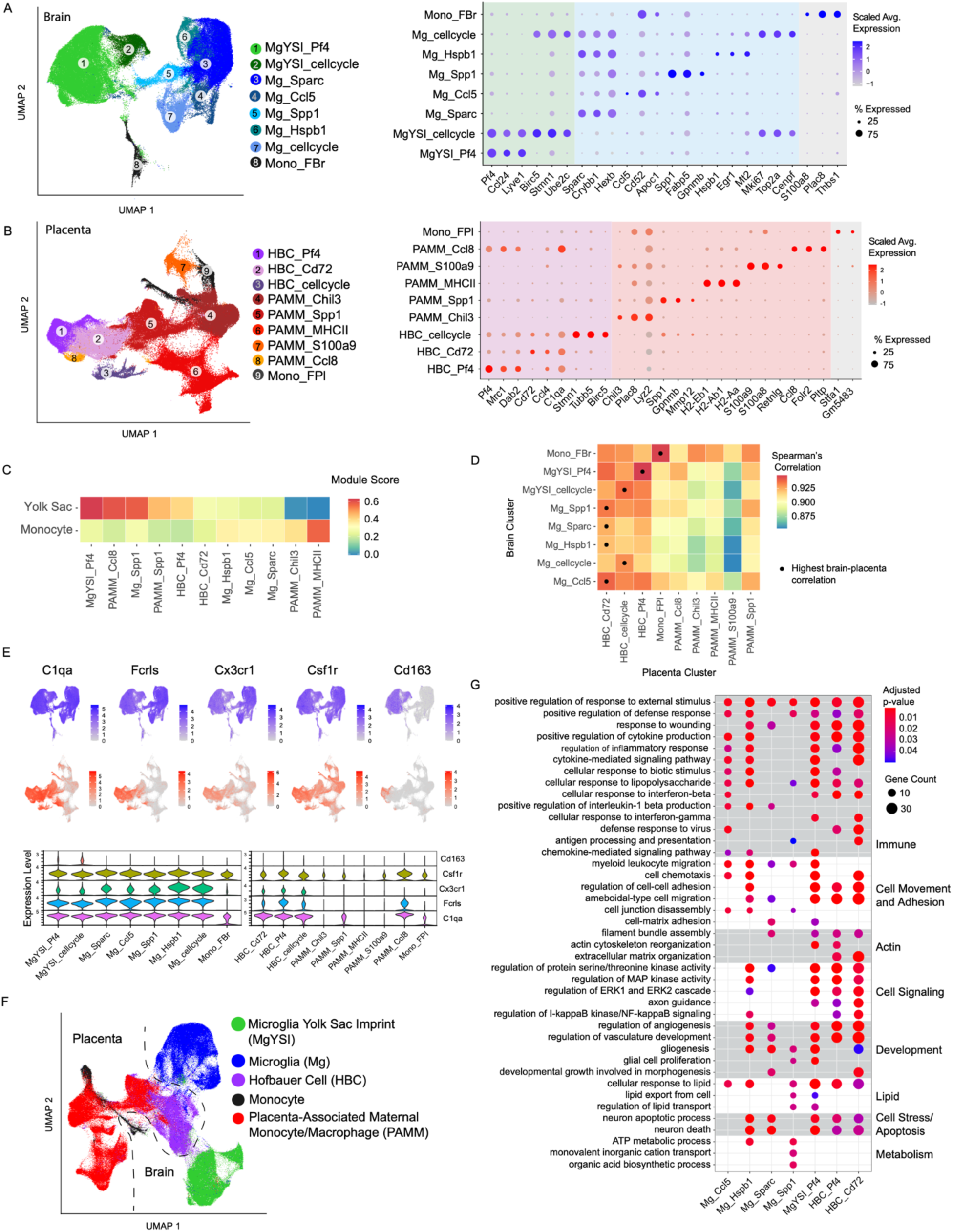
Fetal placental and brain macrophages are heterogeneous populations with shared cluster-specific signatures. **A (left).** Uniform Manifold Approximation and Projection (UMAP) visualization of Cd11b+ macrophage enriched fetal brain microglia/monocyte cells reveals 8 distinct clusters. Unless otherwise specified, clusters are named as “cell-type-prefix_top-marker-gene”. Mg: microglia; Mono_FBr: fetal brain monocytes; YSI: yolk sac imprint. **A (right).** Cluster-average expression of the top 3 marker genes for each cluster (right), dot size indicates the percent of cells expressing the given gene, color intensity represents the scaled average gene expression. **B (left).** UMAP visualization of Cd11b+ macrophage-enriched placenta macrophage/monocyte populations reveals 4 fetal and 6 maternal clusters. Fetal clusters were determined by significantly higher expression of Y chromosome markers *Eif2s3y* and *Ddx3y* relative to expression of X chromosome marker *Xist* (Supplemental Figure 2D-E). Unless otherwise specified, clusters are named as “cell type prefix_top marker gene”. HBC: Hofbauer cell; PAMM: placenta-associated maternal monocyte/macrophages; Mono_FPl: fetal placental monocytes. Color scheme indicates cell origin (purple=fetal macrophages; red=maternal; gray=fetal monocytes). **B (right).** Cluster-averaged gene expression of the top 3 marker genes for each cluster. Dot size indicates the percent of cells expressing the given gene. **C.** Module score for yolk sac-derived macrophages and embryonic liver monocytes^36,38^ **D.** Spearman correlation coefficients of cluster-averaged gene expression between brain and placenta clusters shown in **A, B**. For each brain cluster, the placenta cluster with the highest correlation is shown with a dot. **E.** Expression levels of canonical microglia (top, blue) and Hofbauer cell marker genes (bottom, orange) **F.** UMAP visualization including both brain and placenta clusters shown in **A,B** shows similarity across brain-placenta compartments. **G.** Gene Ontology (GO) Biological Process enrichment analysis for select Microglia and HBC cluster marker genes. The terms displayed here were curated from among the top 25 most significant GO terms, selecting the processes most relevant to macrophage function, and reducing redundancy. Gene Count gives the number of genes in the query set that are annotated by the relevant GO category. GO terms with an adjusted p-value < 0.05 were considered significant. **Full results shown in SI table 2**.

Macrophages at the maternal-fetal interface are known to be a heterogenous population consisting of both maternally and fetally-derived cells. While maternally-derived macrophages at the maternal-fetal interface have classically been referred to as decidual macrophages^30,39,40^, more granular characterization of heterogenous maternally-derived macrophage and monocyte populations has recently been performed^36^, with new nomenclature proposed and adopted: placenta-associated maternal monocyte/macrophages (PAMMs)^36,41^. Distinct subsets of PAMMs were identified in human placenta, including a population of what were previously considered decidual macrophages, as well as subtypes of maternally-derived resident placental macrophages that are distinct from circulating maternal cells^36^. In this work, we characterized PAMMs in murine placenta and examined the similarity between murine and human PAMM signatures.

To distinguish fetal from maternal macrophages, we evaluated expression of male-specific markers DEAD-Box Helicase 3 Y-Linked (*Ddx3y*) and Eukaryotic translation initiation factor 2 subunit 3, Y-linked (*Eif2s3y*) and female-specific marker X-inactive specific transcript (*Xist*) in brain and placenta clusters from male embryos. *Ddx3y*, *Eif2s3y*, and *Xist* were selected for their representative expression after interrogation of a broad X– and Y-chromosome-specific gene expression panel, including the antisense *Xist* transcript X (inactive)-specific transcript, opposite strand (*Tsix*), Ubiquitously Transcribed Tetratricopeptide Repeat Containing, Y-Linked (*Uty),* and lysine demethylase 5D (*Kdm5d)* (Figure S2D-E). Maternally-derived placental macrophage/monocyte clusters were confirmed via high expression of *Xist* and low/no expression of *Ddx3y* and *Eif2s3y* (Figure S2D-E), based on expression in cells isolated from placentas of a male fetus. The same approach applied to brain microglia demonstrated that all cells were fetally-derived (Figure S2D-E), as expected. Using this approach, we identified 3 fetal placental macrophage clusters with transcriptional profiles most closely related to microglia, including HBC_Cd72, HBC_Pf4, and HBC_cellcycle (Figure 2B). These clusters have gene expression profiles similar to human Hofbauer cell clusters described in recent data sets^29,30,36,42^. We also identified one fetal placental monocyte cluster (Mono_FPl), and five PAMM (maternally-derived macrophage) clusters, whose expression profiles were similar to those of recently identified PAMMs from human placenta^36^.

In order to identify the placental macrophage clusters with most similar transcriptional profiles to brain macrophages, we analyzed the correlation of cluster-averaged gene expression (Spearman’s correlation) between all pairs of brain and placental clusters (Figure 2D). These correlation analyses demonstrated that all microglial clusters were more similar to Hofbauer cell clusters than to maternally-derived monocytes and macrophages (PAMMs) or to fetal placental monocytes (Mono_FPl). Multiple microglial clusters closely matched the transcriptional profile of HBC_Cd72, MgYSI Pf4 highly correlated with the transcriptional profile of HBC_Pf4, and both microglial cell cycle signature clusters mapped closely to the HBC_cell cycle cluster. Of the PAMM clusters, PAMM_Spp1 was most closely related to microglial clusters and to Hofbauer cell clusters. Furthermore, combined clustering of both fetal brain and placental macrophages demonstrated that Hofbauer cells are the placental macrophage population with the most similar transcriptional signature to fetal brain microglia (Figure 2F). Both microglia and Hofbauer cells expressed high levels of the canonical tissue-resident macrophage markers *C1qa*, *Fcrls*, *Cx3cr1*, *Csf1r*, and *Cd163* among other (Figure 2E, S2F-G). In sum, both interrogation of the top marker genes by cluster and correlation analysis of cluster-average expression demonstrated the closest relationships between Hofbauer cells and fetal brain microglia, particularly a subset of fetal brain microglia with a strong yolk sac-like signature.

We next sought to understand the conserved biological processes between Hofbauer cells and microglia via Gene Ontology (GO) Biological Process enrichment analyses of cluster marker genes (Figure 2G). Shared processes across populations of Hofbauer cells and microglia can be conceptually grouped into eight broad categories: immune signaling, cell movement and adhesion, actin processes (actin plays a critical role in microglial process elongation, and actin dynamics shape microglial effector functions, such as phagocytosis and response to inflammatory stimuli^43–45^), cell signaling, development (including vascular development/angiogenesis, gliogenesis, hemopoiesis), lipid transport, cellular stress response and apoptosis, and metabolism. The largest HBC cluster, HBC_Cd72, exhibits signatures of immune and inflammatory functions, processes mirrored by microglial clusters Mg_Ccl5, Mg_Hspb1, and Mg_YSI_Pf4. This cluster’s involvement in regulating vascular development, angiogenesis and gliogenesis is most similar to the functional signature of Mg_Hspb1, Mg_Sparc, and Mg_YSI_Pf4. The second largest HBC cluster, HBC_Pf4, is most closely related to Mg_YSI_Pf4 in its signature of actin and lipid-related processes and is engaged in fewer immune, lipid, and protein-related functions in comparison to HBC_Cd72. Both HBC population’s cellular stress response and apoptotic functions are reflective of the functions of all microglial clusters apart from Mg_Ccl5, which is less engaged in these functions. Nearly all microglial clusters and both HBC clusters are engaged in lipid transport. The only microglial functions not closely mirrored by at least one cluster of Hofbauer cells were those related to ATP metabolism and biosynthesis. A complete list of enriched GO biological processes in cluster marker genes may be found in Supplemental Table 2. Taken together, these analyses demonstrate that both fetal brain microglia and fetal placental macrophages are heterogeneous cell types, with Hofbauer cells mirroring microglia in both gene expression and inferred biological processes.

### Fetal placental macrophages provide novel insights into fetal brain microglial programs in the setting of maternal diet-induced obesity

Maternal obesity is associated with increased offspring risk for neurodevelopmental and neuropsychiatric disorders^46–49^, many of which are mediated at least in part by altered microglial functions. Maternal obesity is known to be associated with maternal immune activation^50–53^, which may represent a final common pathway by which adverse microglial programming occurs in the setting of diverse in utero exposures^54–57^. Given the lack of accessibility of fetal brain microglia to interrogate brain immune programs after in utero exposure to obesity or other maternal immune-activating exposures, alternative approaches are needed to accurately understand individual offspring risk. After establishing that Hofbauer cellular programs reflect those of microglia without perturbation, we next sought to examine whether Hofbauer cells could reflect microglial changes in a mouse model of high-fat diet-induced obesity. To understand how placental macrophages and fetal microglia respond to maternal obesity, *C57BL/6J* females were placed on an obesogenic diet (Research Diets D12492, 60% kcal from fat) for 10 weeks to induce maternal obesity (Figure S3A) as described in prior publications by our group^23,58,59^, and we again performed single-cell RNA-sequencing as described above.

To determine the gene expression changes in fetal brain and placental macrophage populations in response to maternal obesity, we performed differential expression analysis within each cell cluster, including fetal brain microglia and placental macrophages from obese compared to control dams, considering male and female fetuses together (Fig 3). 503 unique genes were dysregulated in fetal brain microglia by maternal obesity, and 660 genes in placental macrophages (including 444 in fetal placental macrophages and monocytes and 438 in maternal placental macrophage/monocyte clusters). A complete list of maternal obesity-associated DEG by cluster within fetal microglia and placental macrophages may be found in Supplemental Table 3. In order to characterize the similarity of the response to obesity in microglia and placental macrophages, we compared the DEG in each pair of brain and placental clusters (Fig 3A). We found that genes differentially expressed in the setting of maternal obesity were more often shared between microglial clusters and Hofbauer cells than between microglia and PAMMs; there was a ∼45% overlap between obesity-associated DEG in all microglial clusters and the DEG in HBC_Pf4 and HBC_Cd72, with the exception of microglia cluster Mg_Ccl5, which had only 33% overlap with the HBC DEG. While the Mg_Ccl5 cluster shares with HBC clusters a functional signature related to immunity, cell movement and adhesion, and lipid regulation (Fig 2G), this cluster appears to differ from the other microglial clusters in its lack of marker genes related to actin cytoskeletal functions, cell-cell signaling, angiogenesis/gliogenesis, and functions related to apoptosis and cellular metabolism. Thus overall the Mg_Ccl5 cluster may be engaged in more highly specialized functions than other Mg clusters and thus HBCs may not reflect this cluster as broadly as they do other subpopulations of Mg.

**Figure 3.**
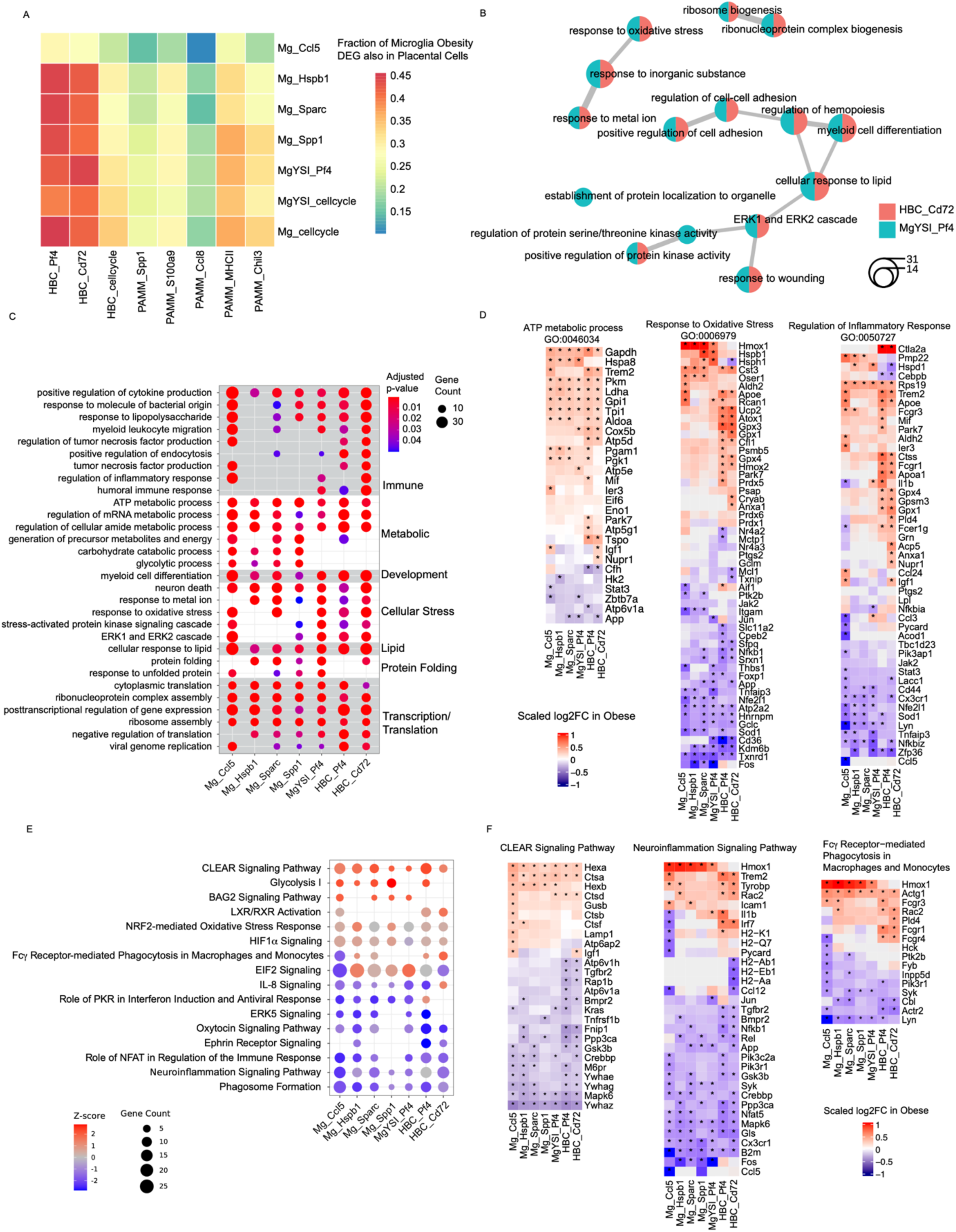
Maternal obesity alters gene expression in fetal microglia and placental macrophages. **A**. Fraction of significantly DEG in obesity-exposed fetal microglia that are also significantly differentially expressed in obesity-exposed placental Hofbauer cells. Differentially expressed genes (DEG) between offspring of obese and control dams are shown. DEG with adjusted p-value <0.05, abs(log2FC)>0.25 are considered significant. **B.** Network plot of the top 15 Gene Ontology Biological Processes enriched in the DEG in fetal MgYSI_Pf4, shown in both MgYSI_Pf4 and placental HBC_Cd72. Nodes correspond to enriched GO categories, node size is proportional to number of genes and edges thickness is proportional to the number of overlapping genes between the two categories. **C.** Enriched Gene Ontology biological processes in the DEG of obesity-exposed microglia and Hofbauer cells. Select results shown, **full results shown in SI Table 4**. Shading corresponds to manual grouping of GO categories. Gene Count gives the number of genes in the query set that are annotated by the relevant GO category. **D.** Up– and downregulation of DEG implicated in ATP metabolism, response to oxidative stress, and regulation of inflammatory response in microglia and Hofbauer cells. DEG fold changes in obesity-exposed microglia and HBCs for three significantly enriched GO biological processes relevant to microglial function.* DEG with adjusted p-value < 0.05. **E.** Select IPA Canonical Pathways enriched in obesity-exposed microglial and Hofbauer cell cluster DEG. Positive z-score means pathway is activated in fetal macrophages of obese dams, and negative z-score means pathway is suppressed. **F.** Up– and downregulation of genes in fetal macrophages of obese dams for two significantly activated or inhibited IPA Canonical Pathways relevant to microglial function. DEG fold changes depicted. * DEG with adjusted p-value < 0.05.

To explore the functional consequences of the gene expression changes induced by maternal obesity, we performed GO enrichment analyses of DEG. Similar to the genes themselves, the GO categories enriched in the DEG of microglial and placental clusters also showed substantial overlap and can be conceptually grouped into six categories: immune signaling, metabolism, development, lipid response, protein folding and transcription/translation (Fig. 3B). In particular, 13 of the top 15 GO biological processes enriched in the DEG of the clusters with the most similar DEG, fetal MgYSI_Pf4 and placental HBC_Cd72, were common to both clusters (Fig. 3C). The majority of the top GO terms were enriched in both clusters, and included processes relevant to microglial function, including Cellular Response to Lipid, Response to Oxidative Stress, and Positive Regulation of Cell Adhesion (Fig. 3C). Consistent with a recent study demonstrating that maternal high-fat diet induces inflammatory responses in the placenta and fetal brain^22^, we noted that immune-related processes were altered throughout Mg and HBC clusters in the setting of maternal obesity. Of note, maternal obesity was associated with a dramatic alteration in metabolic processes in Mg and HBC clusters. The directionality of changes of the genes represented in these processes suggest a shift toward increased energy utilization or glycolysis in obesity-exposed macrophages (e.g., increased *Gapdh*), which is consistent with previous studies suggesting increased glucose metabolism in placentae from women with obesity and Type II diabetes^60,61^. The direction of gene expression changes was overall consistent across Mg and HBC clusters for three key GO categories: ATP Metabolic Process, Response to Oxidative Stress and Regulation of Inflammatory Response (Fig 3D), with a combination of both up– and downregulated genes.

To better understand the direction of effect of functional changes in macrophage populations in response to maternal obesity, we performed Canonical Pathway enrichment analysis using Ingenuity Pathways Analysis (IPA, Qiagen) (Fig 3E). These analyses confirmed a relative activation in glycolysis in both Mg and HBC clusters in the setting of maternal obesity, and identified a relative activation of the Coordinated Lysosomal Expression and Regulation (CLEAR) signaling pathway in Mg and HBC clusters in the setting of maternal obesity. Activation of the CLEAR signaling pathway implies increased lysosomal activity in the setting of increased autophagy secondary to endoplasmic reticulum stress^62–64^. Evaluation of classic macrophage functions such as phagocytosis and inflammatory signaling revealed a complex picture, with both immune activation, but also a macrophage exhaustion phenotype. Specifically, the IPA analysis implicated activation of Fcγ-receptor mediated Phagocytosis in obesity-exposed microglia and HBC clusters, but suppression of Neuroinflammation Signaling Pathway (Fig 3E,F). This mixed picture of both macrophage activation and exhaustion in the setting of maternal immune activation is consistent with and extends published results, which describe not only fetal macrophage priming, activation and increased phagocytosis^22,23^ in response to maternal obesity and other maternal immune-activating exposures, but also macrophage and monocyte exhaustion phenotypes^65,66^.

## Sex differences in fetal brain and placental macrophage responses to maternal obesity

Fetal sex is increasingly recognized as a critical factor influencing placental, fetal and neonatal immune responses ^23,67–69^. Sex differences in the incidence and prevalence of many neurodevelopmental and psychiatric disorders are well established, yet the mechanisms underlying sex differences in these disorders are relatively unknown. Given the critical role of microglia in neurodevelopment, sex differences in microglial development and priming are an attractive candidate mechanism that may underlie sex biases in neurodevelopmental and psychiatric disorders ^70^. Understanding how sex contributes to disease development, susceptibility, and severity is of critical importance when considering clinical implications such as treatment efficacy. Sex differences in the impact of maternal obesity on fetal brain development, microglial and placental function, and neurodevelopmental outcomes have been described^22,53,58,71–73^, but single cell microglial and placental macrophage programs have not yet been investigated in the context of sex differences.

We therefore investigated potential sex differences in the response of fetal placental and brain macrophages to maternal obesity, evaluating DEG in obesity-exposed compared to control fetal brain and placental macrophages in a sex-stratified analysis (Methods, Fig. 4). Despite the fact that cells were isolated on e17.5, prior to the hormonal surge commonly associated with brain masculinization^70^, we observed sex differences in gene expression changes induced by maternal obesity. This suggests that the local microenvironment or other non-hormonal factors are likely driving differences between male and female fetal brain and placental macrophages in late gestation. The number of DEG caused by maternal obesity was significantly higher in all male HBC and PAMM placental clusters compared to female, as well in two microglial clusters, Mg_Spp1 and Mg_Hspb1 (Fig 4A). To facilitate the analysis of sex differences induced by maternal obesity, we combined clusters that are subsets of the same cell type into four cell types: microglia (Mg), yolk sac imprint microglia (MgYSI), HBC and PAMM. Overall, most DEG changed in a similar direction (either upregulated or downregulated by obesity) in both males and females, especially in the two most similar populations MgYSI and HBC (Fig. 4B); we called these genes “sex-consistent” (N= 425 DEG in HBCs and 368 DEG in MgYSI). Genes whose expression changed in opposite directions (e.g. upregulated in males but downregulated in females) in the setting of maternal obesity were designated “sex-dimorphic” (N=58 DEG in HBCs and 92 DEG in MgYSI). GO enrichment analysis was used to determine biological processes that were altered in the maternal obesity-associated DEG in a sex-consistent manner in microglia and HBCs, and pathways that were altered in DEG in a sexually dimorphic manner (Fig. 4B). The GO categories enriched in the sex-consistent genes were similar in microglia and HBC, including categories involving immune response, cell death and protein folding. GO enrichment analysis of the sex-dimorphic genes dysregulated by obesity in both HBC and microglia implicated pathways involved in the regulation of myeloid and leukocyte differentiation and hematopoiesis. Sex-dimorphic genes dysregulated in HBCs alone were concentrated in protein-folding categories. Sex-dimorphic genes dysregulated in microglia alone were implicated in neuronal cell death, response to interferon-alpha, negative regulation of cell-cell adhesion, and actin cytoskeleton-related genes.

**Figure 4.**
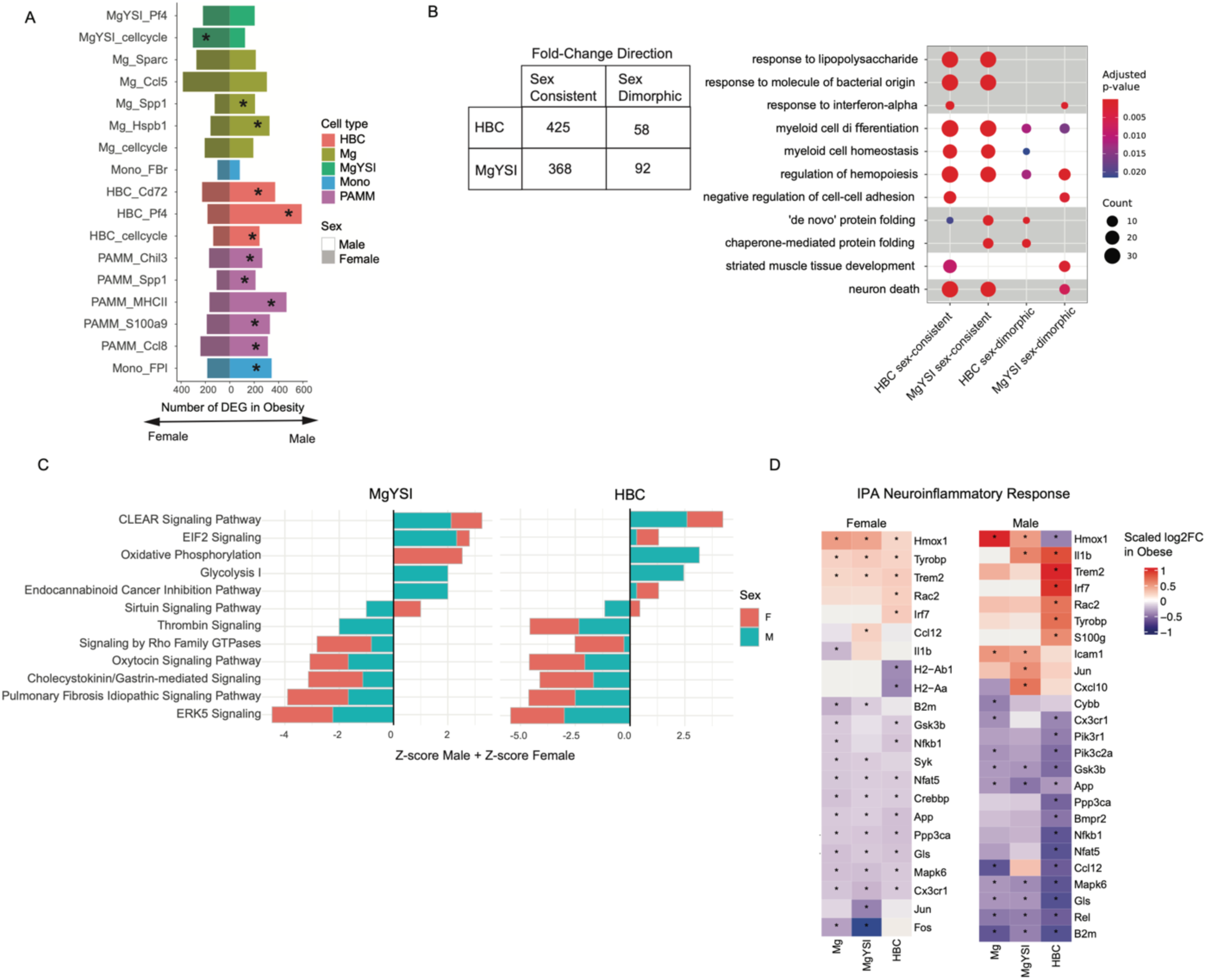
Sex differences in the response of microglia and placental macrophages to maternal diet-induced obesity. Differentially-expressed genes (DEG) between microglia and placental macrophages in fetuses of obese and control dams are shown, calculated separately for male and female offspring. **A.** The number of DEG between obesity-exposed versus control macrophages/monocytes is shown as a bar, colored by cell type, and shaded according to sex. * indicates p<0.05 difference between male and female as given by a 2×2 contingency table Fishers-exact test. **B (left)**. The table considers the total number of DEG in yolk-sac imprint microglia (MgYSI) and HBC cell types, that are DEG in male and/or female cells. Genes are categorized as either having a fold change in the same direction (sex-consistent) or different direction (sex-dimorphic) in males and females. **B (right)**. GO Biological process enrichment results for the set of sex-dimorphic and sex-consistent DEG. **C.** Comparison of the top 12 IPA Canonical Pathways that are enriched in both Mg and HBC. The sum of the Z-score from the female analysis and male analysis is shown, with positive Z-score indicating activation and negative Z-score indicating inhibition. (Full results shown in Supplemental Table 5) **D.** Fold changes in fetal macrophages in offspring of obese versus control dams for the IPA Canonical Pathway NeuroInflammatory Response Pathway for Males and Females.* DEG with adjusted p-value < 0.05.

To determine directionality of pathway activation or suppression, IPA analyses of DEG in males and females were conducted. The activation or suppression of Canonical Pathways in male and female HBCs and MgYSIs were largely similar between sexes (Fig. 4C). In both sexes, maternal obesity was associated with upregulation of pro-apoptotic and pro-autophagy signaling (EIF2 and CLEAR signaling) and downregulation of anti-apoptotic Oxytocin and ERK 5 signaling in both HBCs and MgYSI. In contrast, maternal obesity had a sexually dimorphic effect on sirtuin signaling, with activation in female HBCs and MgYSI and suppression in male HBCs and MgYSI. Increased sirtuin signaling is linked to metabolic control, DNA repair, neuroprotection, and anti-aging effects^74,75^, suggesting a more favorable metabolic response in female HBCs and microglia in the setting of maternal obesity. The upregulation of the Glycolysis I canonical pathway only in male HBCs and MgYSI also suggests less efficient energy utilization by exposed male macrophages compared to female macrophages. In the Neuroinflammatory Response Canonical Pathway, males had a larger magnitude of gene expression fold-changes compared to females, suggesting that male neuroinflammatory response may be more strongly dysregulated in the setting of obesity (Fig. 4D). Fetal sex did not influence cluster identification or the proportion of cells within clusters (Fig S3B,C) with the exception of Mg_Ccl5 and Mono_FBr in male brain, clusters which were slightly more represented in brains of fetuses from control dams (FDR < 0.05).

Taken together, these analyses demonstrated increased numbers of genes dysregulated by maternal obesity in male placental macrophages and two microglial clusters, and stronger dysregulation of neuroinflammatory pathway genes in maternal obesity-exposed male microglia and fetal placental macrophages. These findings extend prior work demonstrating a greater impact of maternal obesity on male versus female fetal brain gene dysregulation^58^ and a greater impact of maternal obesity on pro-inflammatory priming of male fetal brain microglia and placental macrophages^23^. The increased dysregulation of the placental macrophage signature in exposed males relative to the brain macrophage signature is consistent with the known barrier function of the placenta, protecting the downstream fetal organs from varied maternal exposures^76,77^. Maternal obesity was associated with sex differences in the dysregulation of microglial genes related to neuronal cell death, interferon signaling, cell-cell adhesion and motility/phagocytosis. Sex differences in energy utilization and metabolic response to obesity were noted in both fetal placental macrophages and microglia, with female gene expression patterns suggesting a more favorable metabolic response to maternal obesity than male.

## Comparison of Placental Macrophage Signatures Within and Across Species

### Within species comparison: mouse HBCs vs PAMM

In order to employ HBCs as a useful cell population to evaluate fetal neuroimmune development, it is important to distinguish them from other placental macrophages with distinct developmental trajectories (e.g., PAMMs). Through sequencing placental macrophages at single-cell resolution, our study has characterized heterogeneity in HBC and PAMM populations at unprecedented resolution. Marker genes that were more highly expressed in HBC compared to PAMM included *Complement 1q c chain* (*C1qc*), *Complement 1q a chain* (*C1qa*), *Platelet factor 4* (*Pf4*), *C-C motif chemokine ligand 4 (Ccl4*), *DAB adaptor protein 2* (*Dab2*), and *Mannose receptor c type 1* (*Mrc1*), whereas *Placenta associated 8* (*Plac8*) as well as several MHC-II genes were more highly expressed in PAMMs (Fig. 5A). Relative to PAMM markers, HBC marker genes were specifically enriched for GO biological processes related to chemokine signaling, ERK1/2 signaling, and interestingly, regulation of neuron death (highlighting the potential for HBCs to provide information about microglial-mediated neuronal apoptosis^78,79^). PAMM markers were specifically enriched for MHC-II-mediated antigen presentation and processing, regulation of T-cell activation, and cell-cell adhesion processes (Fig. 5B). In contrast with recent observations in human first trimester placenta^36^, *Folr2* was not as strong a marker for mouse HBCs (Fig. 5C,D), but other key HBC (*Mrc1*) and PAMM (MHC-II gene *H2-Eb1*, *S100a9*) markers from the same study were well-represented in mouse HBC and PAMM subsets. These data suggest that while mouse and human placental macrophages have similar signatures, Folr2 in particular may not be the strongest marker for HBC in mice, at least at the end of gestation. Furthermore, we calculated the marker genes with valid antibodies that would best distinguish HBC from PAMM (Fig. 5 E), including positive selection markers Mrc1 and Cd83, and negative selection markers Thbs1 (primarily enriched in placental monocytes and PAMM) and Cd74 (primarily enriched in PAMM).

**Figure 5.**
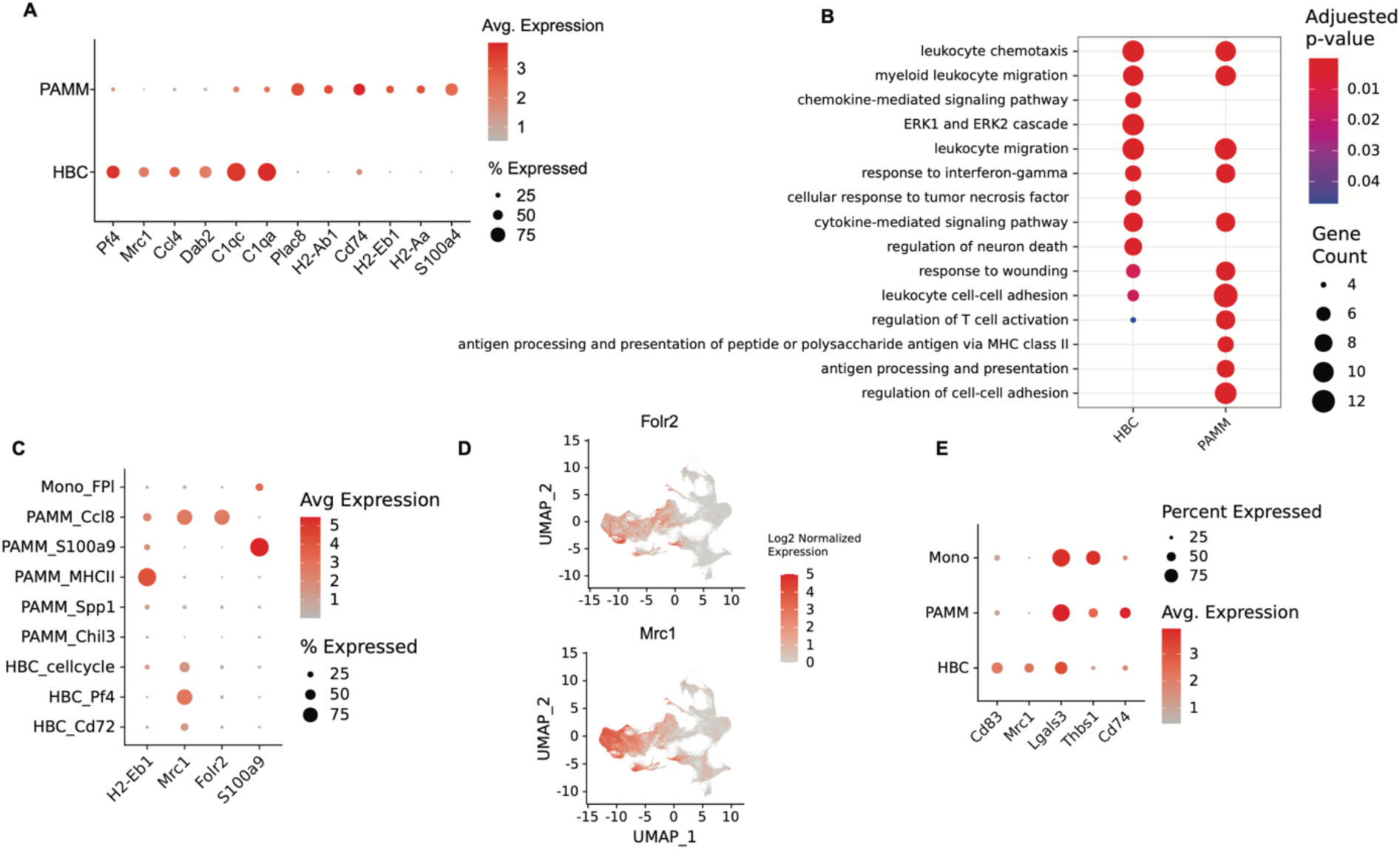
Comparison of maternal and fetal placental macrophages. **A**. Cluster-average expression of top marker genes distinguishing PAMM from HBC. Dot size indicates the percent of cells expressing the given gene. **B.** Gene Ontology (GO) Biological Process enrichment analysis for for HBC and PAMM marker genes. Gene Count gives the number of genes in the query set that are annotated by the relevant GO category. GO terms with an adjusted p-value < 0.05 were considered significant. **C.** Cluster-average expression of genes that distinguish PAMM and HBC populations in human studies. **D.** Detailed expression of genes Folr2 and Mrc1 that distinguish PAMM and HBC populations in Thomas 2020.^36^ **E.** Markers with valid antibodies that best separate HBC, placental monocyte and PAMM populations.

### Across species comparison: mouse vs. human HBCs and PAMM

To build upon the observation that *Mrc1, Cxcr3, Cd83* and *Adgre1*, but not *Folr2*, are the best surface markers by which to positively select HBCs in mouse placenta, we sought to understand the extent to which mouse placental macrophages are similar to human placental macrophages. Mouse microglia are widely accepted as an excellent model for human microglia ^16,80,81^, but substantially less is known about the ability of mouse placental macrophages to model their human equivalent. Mice and humans are known to share key similarities in immune system development, including the yolk sac as the initial site of hematopoiesis and origin for granulo-macrophage progenitors (e.g. the CX3CR1+ cells that give rise to microglia and Hofbauer cells)^82–86^ and conserved cell surface antigen expression in yolk sac-derived macrophages ^16,87^.

While the presence of resident fetal macrophages in the mouse placenta has been documented by our group and others^23,88–90^, the lineage tracing experiments detailed earlier in the manuscript help confirm the definitive yolk sac origin of a majority of these cells. Despite this knowledge, mouse HBC have never been profiled using scRNA-seq, and these data have never been compared to those from human placentas. We therefore compared our mouse scRNA-seq data to several reference human datasets that contained high representation of HBC, representing human placentas obtained from 6 weeks through 40 weeks (full term)^29,30,42^. Comparison of our scRNA-seq data to scRNA-seq data from human placenta datasets demonstrated strong correlation of cluster-averaged gene expression between mouse Hofbauer cells and human clusters identified as Hofbauer cells (notated as “villous Hofbauer cells, vil. HC, HB or vil.Hofb”) at 6-11 weeks ^29^ (Fig S4A), 6-14 weeks ^30^ (Fig S4B), and full-term ^31^ (Fig S4C), with Spearman Correlation 0.71-0.76. Our murine Hofbauer cell and PAMM clusters also mapped closely to human Hofbauer cells in these human single cell datasets. Taken together, these data demonstrate that placental resident macrophage and monocyte single-cell gene expression signatures are highly conserved across mice and humans.

## Discussion

Microglia, brain resident macrophages, are vulnerable to diverse maternal immune-activating exposures including bacterial and viral infection, metabolic inflammation such as that mediated by obesity and diabetes, environmental toxicants, and maternal stress, potentially contributing to adverse neurodevelopmental and psychiatric outcomes in offspring^10,46,53,91–96^. However, the mechanisms that dictate susceptibility to these outcomes are unclear, and there is a critical need to identify offspring at greatest risk. To identify more accessible readouts of aberrant microglial priming conferring risk for neurodevelopmental and psychiatric disorders ^10–13^, we sought to evaluate whether extra-embryonic placental macrophages have similar transcriptional signatures as fetal brain microglia at baseline, and if placental macrophages can provide information about the impact of a maternal perturbation, diet-induced obesity, on fetal brain immune programs. Here, we demonstrate (1) a common yolk sac origin for microglia and Hofbauer cells, and (2) similar transcriptional programs between subsets of microglia and Hofbauer cells both at baseline and (3) in response to maternal diet-induced obesity, providing evidence that term placental Hofbauer cells have the potential to uniquely inform on the programs of fetal microglia. We further demonstrate (4) sex differences in the impact of maternal obesity on fetal brain and placental macrophages, including more genes dysregulated in male placental macrophages and two male microglial clusters, and stronger dysregulation of neuroinflammatory genes in the male brain. Female placental and brain macrophages appeared to demonstrate a more favorable metabolic response than male macrophages in the setting of maternal obesity, with upregulation of sirtuin signaling in females and downregulation in males, and upregulation of glycolysis in male macrophages only. Finally, we show (5) significant correlations between murine and human placental macrophage populations, suggesting the potential for Hofbauer cells to provide a readout of neurodevelopmental exposures in humans, as well. The shared ontogeny and transcriptional programs between fetal placental macrophages and fetal brain microglia indicate that term Hofbauer cells, readily accessible at birth, may provide insight into microglial state and function after specific in utero exposures.

Previous functional studies have suggested that the immune functions of Hofbauer cells and microglia are strongly correlated^23,97^, including prior work by our group demonstrating differential cytokine response following TLR4 stimulation in the setting of maternal obesity^23^. However, this prior work left multiple unanswered questions about whether Hofbauer cells might be able to serve as a proxy for microglial cells. To our knowledge, no previous studies have investigated whether murine or human Hofbauer cells reflect microglia transcriptionally, particularly in the setting of maternal obesity. In addition to similarities at baseline, we sought to establish whether these cells would reflect microglia in their response to a maternal exposure such as obesity. In addition, while maternal obesity is known to be associated with adverse neurodevelopmental outcomes in offspring that may be more pronounced in males^53,71^, the impact of maternal obesity on microglial and placental macrophage programming, sex differences in this regard, and how to best identify vulnerable offspring, had not yet been described.

Consistent with prior studies, we observed significant microglial heterogeneity in fetal brains^34,35,98^. Not surprisingly, the microglial clusters with the more robust yolk sac signature also highly express genes known to be associated with another yolk sac-derived tissue resident brain macrophage population, CNS-associated macrophages (CAMs)^99–101^. Our results also extend prior work in understanding fetal microglial responses to maternal obesity. We observed a mixture of activation and suppression of immune related processes in fetal microglia and placental macrophages in response to maternal obesity. Our lab previously reported that obesity primed Mg and HBCs toward a pro-inflammatory phenotype when treated with LPS^23^. More recent studies show that macrophage and monocyte exhaustion plays a role in obesity-associated cellular programming^66^.

Prior work has demonstrated sex-specific fetal brain gene expression signatures using whole fetal forebrain in a murine model, without subselection for any specific cell or immune cell subtype^58^. Given a strong sex bias in many microglial-mediated neurodevelopmental disorders, with autism spectrum disorder, attention deficit hyperactivity disorder, and cognitive delay/learning disabilities all more common in males than females ^53,70,102,103^, and the fact that male fetal macrophages (both brain and placental) may be more vulnerable to pro-inflammatory intrauterine priming compared to female macrophages ^23,97^, we sought to investigate potential sex differences in fetal microglia and placental macrophage responses to maternal obesity. Our finding that significantly more genes were differentially regulated by obesity in male compared to female placental macrophages, is consistent with prior work demonstrating a disproportionate impact of maternal obesity on male versus female placental immune dysregulation^22,23,104^. The significantly greater number of DEG by maternal obesity in the male placenta compared to the male fetal brain is consistent with the known barrier role played by placenta to shield downstream fetal development from various maternal exposures^105,106^. The male fetus may be more vulnerable to maternal exposures, mediated in part through increased male placental reactivity to maternal exposures^69,107–109^.

Beyond the specific similarities between Hofbauer cells and microglia, we identified two elements that should facilitate the application of Hofbauer cells as proxies for microglia in future murine and human studies. First, despite their recognized importance in immune signaling at the maternal-fetal interface ^110^, Hofbauer cells have been under-characterized due to challenges in cell isolation and maintenance of viability ^111^. While microglia have been demonstrated to be a highly heterogeneous cell type with subsets of cells performing specialized functions ^34,35^, there is a relative lack of knowledge regarding Hofbauer cell subsets and their functions. By enriching for macrophages and monocytes with a Percoll gradient followed by sub-selection for CD11b+ cells, our experiments were able to provide uniquely targeted single-cell sequencing and greater resolution on placental macrophage populations than has been previously achieved. This increased resolution yielded new transcriptional and functional insights about the under-characterized heterogeneity of Hofbauer cells and PAMMs. Consistent with other groups ^30,36,112^, we found that maternal cells contribute significantly to placental macrophage population. Our use of male placentas and a panel of X– and Y-chromosome markers to identify fetal versus maternal placental macrophages represents an advance in understanding, permitting more precise characterization of the transcriptional profile and function of fetal versus maternal placental macrophages.

The identification of markers more specific to Hofbauer cells should also enable their investigation as biomarkers by more readily distinguishing them from PAMMs. Our cell isolation protocol was selected due to its ability to permit isolation of fetal brain and placenta macrophages in parallel, in a timely fashion to optimize cell viability for sequencing. In our study with multiple male and female embryos, we could use Y-chromosome specific markers in male samples to resolve maternal from fetal macrophages in the placenta, and extrapolate those insights to placental clustering in both sexes. One necessary step to extend these results to be used in clinical practice is to identify Hofbauer cell surface markers that clearly differentiate them from other cell types. While Folr2 demonstrated promise as such a marker for Hofbauer cells in first trimester human placenta ^36^, in late gestation murine Hofbauer cells, we show for the first time that Mrc1 is a better candidate HBC marker, and specific MHC-II cell surface markers (H2-Eb1, H2-Aa, H2-Ab1) may be used to distinguish PAMMs.

As a further step toward application of these cells to study human disease processes, we found strong evidence that murine Hofbauer cells can be used to model human Hofbauer cells. Parallels between human and murine placentation are apparent in both gross morphology and function as well as molecular pathways^82–86^. Although the mouse placenta does have certain fundamental differences from human placenta, such as the degree of contact between fetal tissues and maternal blood, or variability in tissue structure between human placental villi and mouse fetal placental labyrinth, both species share important functions in nutrient transport, blood filtration, and immunocompetency ^37,113,114^. Key similarities between mouse and human immune system development relevant to these experiments include the yolk sac as the initial site of hematopoiesis and origin for both microglia and Hofbauer cells^82–86^ but certainly differences remain between human and mouse immune development ^115^. We interrogated the transcriptional relationships between our Hofbauer cell and PAMM clusters and those of three independently published human data sets^29,30,42^. In all comparisons, the signatures of murine Hofbauer cells and PAMMs from our dataset were closely correlated with human Hofbauer cell and PAMM signatures, suggesting a highly conserved evolutionary relationship. Taken together, these findings provide strong evidence that the mouse placenta can be used to understand immune interactions at the maternal-fetal interface.

In summary, these data provide a precedent for using placental Hofbauer cells as a noninvasive biomarker of fetal brain microglial programming, without perturbation as well as in the setting of an in-utero exposure to maternal diet-induced obesity. These results extend previous work from our groups and others demonstrating that Hofbauer cells and fetal brain microglia respond similarly to bacterial infection ^97^, and show similar exaggerated responses to the bacterial endotoxin LPS after priming by maternal obesity ^23^. They lay the groundwork for longer-term translational studies in humans correlating Hofbauer cell inflammatory profiles with offspring neurological outcomes. Such studies will determine whether Hofbauer cells can serve as a clinical indicator of neurodevelopmental vulnerability, with the ultimate goal of identifying vulnerable offspring at birth and facilitating interventions during key developmental windows of plasticity.

## STAR Methods

### Resource Availability

#### Lead contact

Further information and requests for resources/reagents should be directed to and will be fulfilled by the lead contact, Andrea Edlow

#### Materials availability

This study did not generate new unique reagents

#### Data and code availability

Single-cell RNA-seq data will be deposited in GEO and will be publicly available as of the date of publication. Single-cell RNA-seq count matrices are available at: https://data.mendeley.com/preview/r422vr5gbm?a=f2f15f2b-e3c0-40dd-949d-c4b821babbd4. Accession numbers are listed in the key resources table. All code necessary to reproduce these analyses is available at https://github.com/rbatorsky/fetal-mac-edlow. Any additional information required to reanalyze the data reported in this paper is available from the lead contact upon request.

## Experimental Model and Subject Details

### Animals

#### Strains and husbandry conditions

All procedures relating to animal care and treatment conformed to Massachusetts General Hospital Center for Comparative Medicine Program, Duke University Animal Care and Use Program, and NIH guidelines. Animals were group housed in a standard 12:12 light-dark cycle. The following mouse lines were used in this study: *FVB-Tg(Csf1r-cre/Esr1*)1Jwp/J* (Jackson Laboratory, stock no. 019098, referred to as *Csf1r-Cre^ER^*hereafter), *B6.Cg-Gt(ROSA)26Sor^tm^*^14^*^(CAG-tdTomato)Hze^/J* (Jackson Laboratory, stock no. 007914, referred to as *tdTomato^f/f^*hereafter), and *C57BL/6J* (Jackson Laboratory, stock no. 000664). *Csf1r-Cre^ER^* animals were backcrossed to *C57BL/6J* mice for one generation prior to breeding with *TdTomato^f/f^* animals. *Csf1r-Cre^ER^* was maintained in the males for all experimental studies, and genotyping was performed per Jackson Laboratory published protocols for each strain. For diet-induced obesity versus lean control experiments, *C57BL/6J* females were placed on either an obesogenic diet (Research Diets D12492, 60% kcal from fat) for 10 weeks to induce maternal obesity, or a control diet (D12450J, 10% fat) matched for protein, fiber and sucrose content for the same duration of time, prior to breeding with *C57BL/6J* males on the control diet, as described in prior publications by our group^23,58,59^. Pre-breeding dam weight curves and gestational weight curves are depicted in Figure S3A. Pregnant *C57BL/6J* dams were euthanized at gd17.5 and embryonic brains and matched placentas were retrieved as described below (Tissue collection for sequencing).

#### Transgenic Breeding/Maintenance

Male *Csf1r-Cre^ER^;TdTomato^f/f^* mice were crossed with *TdTomato^f/f^* or *Tdtomato^f/+^* females to generate control *TdTomato^f/f^* or *Tdtomato^f/+^* and experimental *Csf1r-Cre^ER^;TdTomato^f/f^* or *Csf1r-Cre^ER^;TdTomato^f/+^* animals within the same litters. We did not see any spontaneous recombination (e.g. tdTomato fluorescence or RFP immunoreactivity) in control animals. Pregnancy was determined by the presence of a copulation plug (gestational day 0.5 (gd0.5)), and maternal weight was measured at gd0.5 and gd8.5 (to confirm pregnancy weight gain). Pregnant females were injected intraperitoneally (i.p.) with 10 mg/kg 4-hydroxytamoxifen (4-OHT, Millipore-Sigma cat #H6278) dissolved in corn oil (Sigma) at gd8.5 and euthanized at gd17.5 with CO_2_ followed by rapid decapitation. Sex was recorded and is clearly noted throughout the manuscript.

#### Tissue collection for immunohistochemistry

Uterine horns containing embryos were rapidly dissected and placed on ice in sterile 1X PBS. Individual embryos were separated, and placenta, brain, and tail tissue (for genotyping) were collected. Placenta and brain tissue were fixed in 4% paraformaldehyde in PBS (PFA, Sigma) overnight at 4°C, cryoprotected in 30% sucrose + 0.1% sodium azide in PBS (Sigma), and embedded in OCT (Sakura Finetek, Torrance, CA) before being cryo-sectioned. Sections were frozen at –80°C for storage. 40µm cryosections were collected directly onto Superfrost slides (Fisher), permeabilized in 1% Triton X-100 in PBS, and blocked for 1 hr at room temperature using 5% goat serum (GS) in PBS + 0.1% Tween-20. Sections were then incubated for 2 nights at 4°C with chicken anti-Iba1 (Synaptic Systems, 234 006) and rabbit anti-RFP (Rockland, 600-401-379). Following PBS washes, sections were then incubated with anti-rabbit Alexa-594 (placenta and brain), anti-chicken Alexa-647 (placenta) or anti-chicken Alexa-488 (brain) (1:200; ThermoFisher), and DAPI (100µg/mL). Ten z-stacks of 0.5µm optical thickness were taken using a Zeiss AxioImager.M2 (with ApoTome.2) from at least 5 sections (each 400µm apart) from the fetal compartment of each placenta. Ten z-stacks of 0.5µm optical thickness were also taken from at least 3 hippocampal sections (each section being 200µm apart).

#### Tissue collection for sequencing

Uterine horns containing embryos were rapidly dissected and embryos were placed on ice in sterile PBS. Brains and placentas were isolated, and fetal forebrain and placenta were diced finely with sterile blade and placed into collagenase A (Millipore-Sigma; 11088793001) digestion solution on ice. Cd11b-positive cells were isolated from both placenta and brain tissue as previously described^23,116^. Briefly, samples were serially dissociated into a single cell suspension using hand-flamed Pasteur pipettes, and the resultant suspension was enriched for macrophages and monocytes using a 70%/30% Percoll gradient (Millipore-Sigma; GE17-0891-01). This enriched macrophage/monocyte solution was incubated with human and mouse CD11b microbeads (Miltenyi Biotec; 130-093-634), and cells were further enriched for Cd11b-positive macrophages/monocytes using Miltenyi MACS Separation Columns (LS; 130-042-401) and a Miltenyi QuadroMACS Separator (130-090-976). Fresh CD11b+ cells from 8-9 biological replicates per diet group (4 male fetal brains and matched placentas per diet group, 4-5 female fetal brains and matched placentas per diet group) were then prepared for single-cell RNA sequencing (10X Genomics v3.0 chip). Paired fetal brain and placental samples were collected for 17 mice, giving 34 total samples. Approximately 5,900 cells/sample were sequenced, giving 197,000 sequenced cells with an average depth of approximately 22,000 reads/cell).

### Quantification and Statistical Analysis

#### Immunohistochemistry quantification

Maximum intensity projections were generated with FIJI (FIJI Is Just ImageJ^117^), and the Cell Counter plugin was used to aid in manual counting of Iba1+ and tdTomato/RFP+ cells. Counts were done by an individual blinded to the age, sex, and genotype of the tissue. Sample sizes can be found in the legend for Figure 1.

#### Single cell RNA-sequencing analysis

10× Data analysis and clustering: 10× scRNA-seq data was aligned with 10× Genomics Cell Ranger (v3.0)^118^ against the mm10 mouse reference genome. Downstream analysis was performed with R (v4.0.0) package Seurat (v4.3.0)^119–122^. Initial filtering was performed on each sample as follows: cells with UMI count <500, gene count <250, or fraction of reads aligning to mitochondrial genes > 0.2 were removed; genes detected in < 10 cells were also removed. All samples were normalized and putative doublet cells were removed using predictions from DoubletFinder (v2.0.3)^123^. All samples were integrated to remove batch effects from individual animals using the Seurat Single Cell Transform workflow ^124^ using the top 3000 variable features, followed by clustering using the Louvain algorithm on the shared nearest neighbor graph, as implemented in the Seurat FindClusters function with resolution 0.4, and visualization by UMAP using the first 50 dimensions. Cluster marker genes were identified using Seurat function FindAllMarkers. Markers were considered significant with adjusted p-value < 0.05 and average log fold-change between two groups > 0.25.

#### Identification of cell types

Annotation of clusters with cell types was done in three steps: 1) First, clusters were annotated using R package SingleR (v1.2.4) with celldex (v.0.99.1) ^125^ packages built-in MouseRNAseq reference; 2) annotation was confirmed using the top Spearman correlation coefficient between cluster-averaged gene expression in our clusters and cluster-averaged gene expression cell types from published datasets for mouse microglia^33–35^ and human placenta^29,30,42^ using the cluster correlation method recently described^31^ (results shown in Figure SI4); 3) cell-type assignment was refined by manual examination of marker genes. Clusters identified as Monocyte– or Macrophage-like were selected and re-clustered (the full set of identified clusters is shown in Figure S2B-C). Single-cell gene signature enrichment scores were calculated using the AddModuleScore function in Seurat using gene signatures described by Thomas et al ^36^. The test for differential cell-type proportions was conducted using a moderated t-test on the logit-transformed proportions as implemented in the propellor function from the R package Speckle^126^. Marker genes with valid antibodies were selected using the Detect_single_marker function from the R package sc2marker^127^.

#### Differential gene expression analysis

Differentially expressed genes (DEG) were calculated with each cluster or cell-type in fetal brain and placental cells between obese and control dams using the FindMarkers function in Seurat with arguments test.use=”MAST” and latent.vars=”sample”. Genes were considered significantly differentially expressed with adjusted p-value < 0.05 and absolute log2 fold change > 0.25.

#### Functional enrichment analyses

Gene Ontology Biological Process enrichment analysis was performed on both cluster marker genes and differentially expressed genes using R package clusterProfler (v. 4.0.5)^128^ and underlying database AnnotationDb org.Mm.eg.db (v3.13.0). IPA Canonical Pathway and Diseases and Functions analysis performed with IPA (Content Version: 90348151). GO terms and IPA pathways with adjusted p-value < 0.05 were considered significantly enriched.

## Supplemental Files

**Supplemental Table 1.** Complete list of cluster marker genes for brain and placental cell-types shown in Fig. 2. avg_log2FC = log base 2 fold-change of the average expression between the two groups. Positive values indicate that the gene is more highly expressed in the first group pct.1 The percentage of cells where the gene is detected in the first group pct.2 The percentage of cells where the gene is detected in the second group p_val_adj Adjusted p-value, based on Bonferroni correction using all genes in the dataset.

**Supplemental Table 2.** GO Biological Process enrichment results for cluster marker genes from brain and placenta cell-types shown in Fig. 2. GeneRatio: Ratio of the size of the overlap of the input geneset with the GO category of interest to the size of the overlap of the input geneset with all the members of the collection of GO categories. BgRatio: Ratio of the size of the input gene set to the size of all of the unique genes in the GO database. Count: size of the overlap of the input geneset with the GO category of interest.

**Supplemental Table 3.** Complete list of differentially expressed genes by maternal obesity for a) Males and females combined and b) Sex stratified. Columns as in SI Table 1.

**Supplemental Table 4.** GO biological process enrichment for maternal obesity-associated DEG for a) Males and females combined and b) Sex stratified. Columns as in SI Table 2.

**Supplemental Table 5.** IPA Canonical Pathways analysis results for maternal obesity-associated DEG for a) Males and females combined and b) Sex stratified. Ratio – the number of molecules in a given pathway that meet your cutoff criteria divided by the total number of molecules that make up that pathway and that are in the reference set. z-score – predicted activation state.

**Supplemental Table 6.** HBC vs. PAMM a) marker genes and b) GO biological process enrichment. Columns as in SI Tables 1,2.

## Author Contributions

R.B. and A.M.C. contributed equally, and as co-first authors. A.G.E. conceived the study and, together with S.D.B., A.M.C., and R.B., designed the experiments. Acquisition of data: A.M.C., R.B., S.K., E.A.B., L.L.S., A.G.E. Analysis and interpretation of data: R.B., A.M.C., B.A.D, D.K.S., S.D.B, A.G.E. Drafting of the manuscript: R.B., A.G.E., A.M.C. Revising the manuscript critically for important intellectual content: A.M.C., R.B., L.L.S., S.K., E.A.B., B.A.D., R.H.P., D.K.S., S.D.B., A.G.E. All authors have given final approval for submission.

## Funding

Research reported in this publication was supported by the Eunice Kennedy Shriver National Institute of Child Health & Human Development (R01 HD 100022 to A.G.E., F32HD104430 to A.M.C, 5K12HD103096-04 to L.L.S.), the Robert and Donna Landreth Family Foundation, and the Charles Lafitte Foundation (awards to SDB).

## Declaration of Interests

A.G.E. serves as a consultant for Mirvie, Inc. outside of this work. A.G.E. receives research funding from Merck Pharmaceuticals outside of this work. R.H.P. is a founder and member of the scientific advisory board of Psy Therapeutics; a member of scientific advisory boards for Swan AI Studio, Belle Artificial Intelligence, Genomind, and Circular Genomics; and consults to Alkermes, Burrage Capital, and Vault Health. He serves as an associate editor for JAMA Network Open. All of these roles are outside the present work.

## Inclusion and Diversity

We support inclusive, diverse and equitable conduct of research.

## Acknowledgements

We thank Dr. Abby Groff and Dr. David Page for advice regarding identification of male versus female macrophages.

