## Supplemental Figures for "Hofbauer cells and fetal brain microglia share transcriptional profiles and responses to maternal diet-induced obesity"

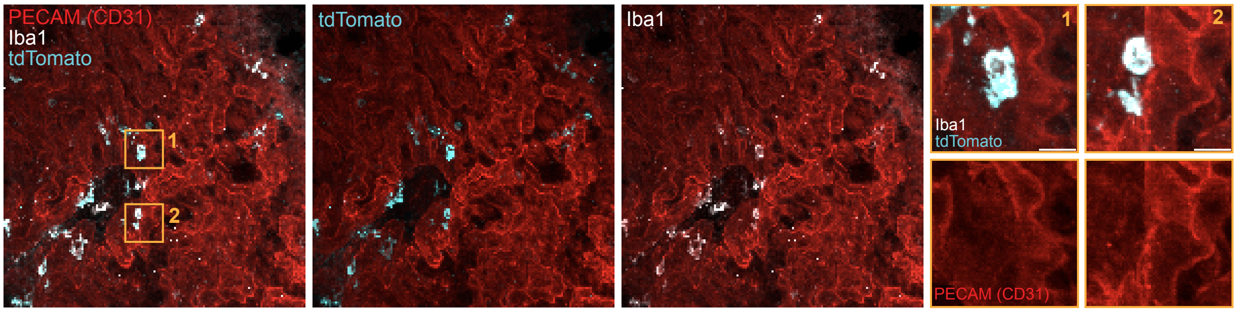


**Supplemental Figure 1: Fetal placental macrophages are parenchymal and not restricted to the vasculature.**
Immunohistochemistry for endothelial cells (PECAM [CD31]), the macrophage marker Iba1, and fetal yolk sac derived macrophages (Csf1R-CreER;TdTomatof/f reporter) demonstrates that fetal placental macrophages are not restricted to the vasculature at e17.5. Scale bar = 50 µm, Inset scale bar = 10 µm

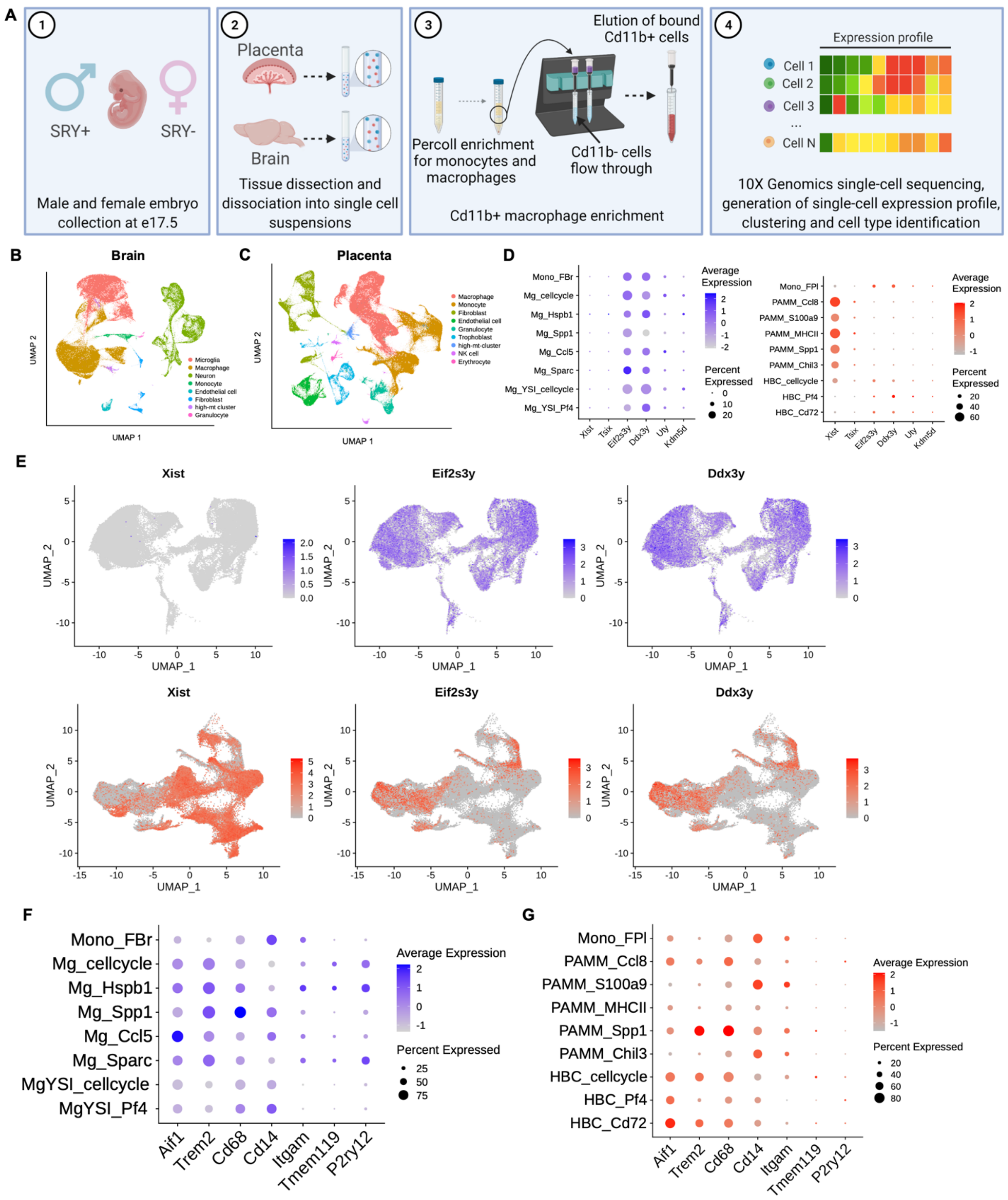


**Supplemental Figure 2: Experimental strategy and identification of fetally- versus maternally-derived macrophage clusters**

**A.** Schematic depicting experimental paradigm. This figure was made using Biorender.

**B.** Uniform manifold Approximation and Projection (UMAP) plots of all fetal brain cell-type clusters.

**C.** UMAP of all placental cell-type clusters.

**D.** Dot plot of all male-specific marker expression in (left) fetal brain cell types and (right) placenta cell-types.

**E.** UMAPs of representative female-specific (*Xist*) and male-specific (*Eif2s3y* and *Ddx3y*) marker expression in (top) male fetal brains and (bottom) placentas from male fetuses.

**F.** Dot plot depicting canonical microglia marker expression in (left) fetal brains and (right) placental macrophages.

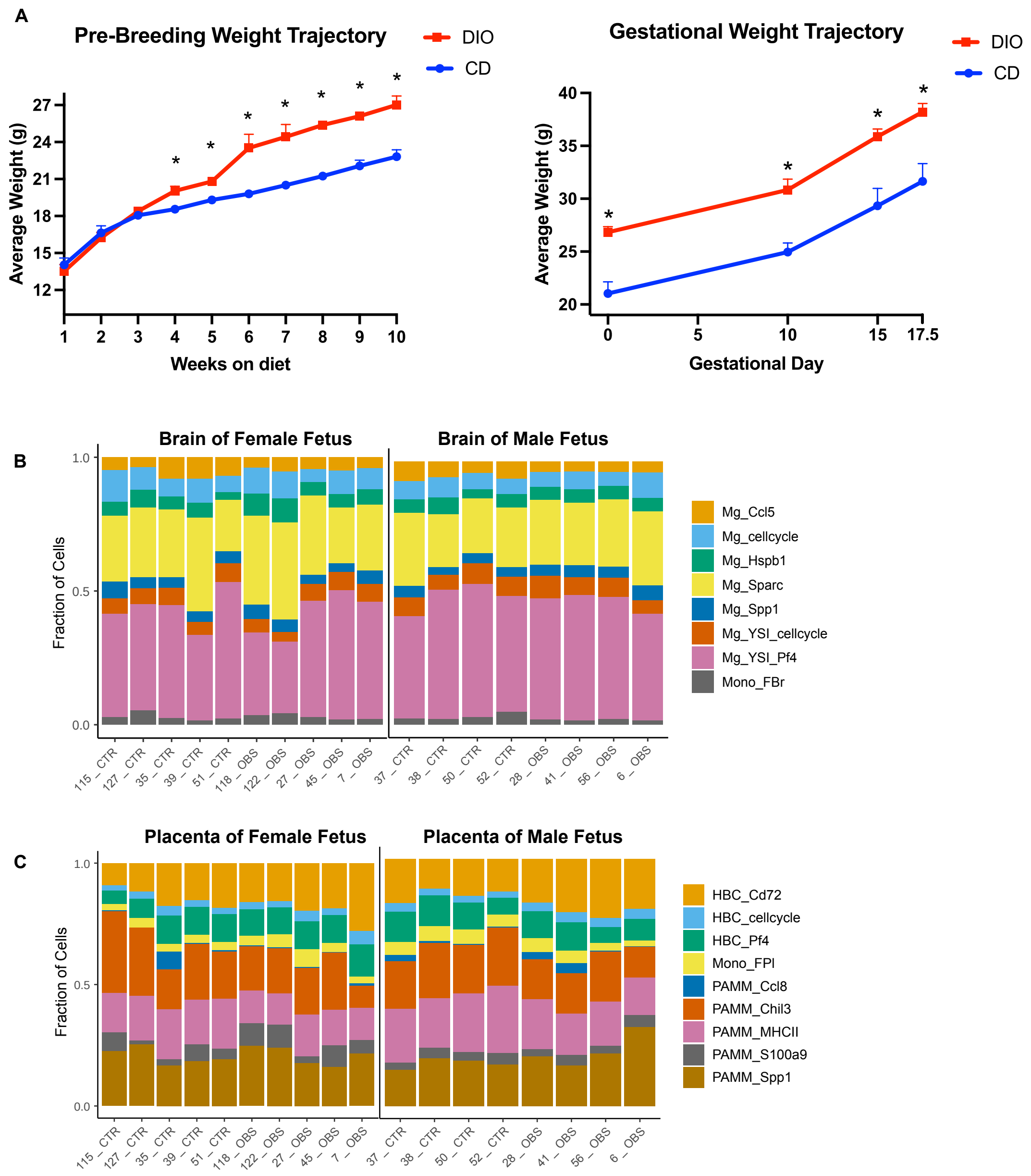


**Supplemental Figure 3: Diet-induced obese and control dam weight trajectories and representation of individual fetuses across microglial and placental macrophage cell clusters**

**A. (left)** Pre-breeding weight curves and **(right)** Gestational weight trajectory for diet-induced obese (DIO) and control dams.

**B.** Composition of fetal brain cells from obese and control dams. Stacked bar plots showing the proportion of each cell group in each sample. Fraction of total cells made up by each annotated cell type is plotted on the Y axis.

**C.** Fractions of placental cells from obese and control dams. Stacked bar plots showing the proportion of each cell group in each sample. Fraction of total cells made up by each annotated cell type is plotted on the Y axis. Test for significance of different cell-type proportions in obese vs. control (Methods) showed Mg_Ccl5 and Mono_F_Br were more prevalent in the brains of male fetuses from control dams (FDR < 0.05, data not shown).

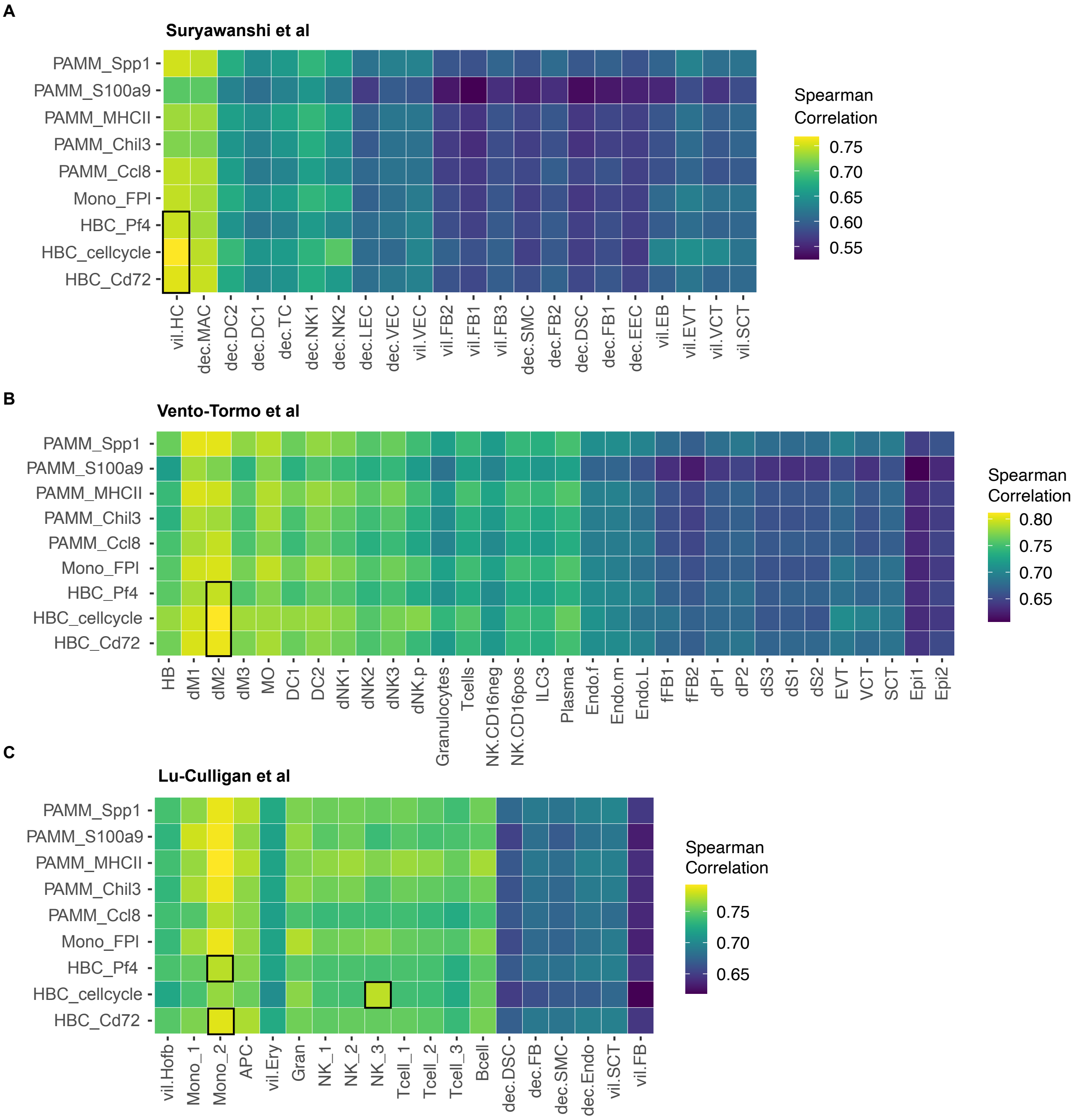


**Supplemental Figure 4**: **Correlation between murine and human placental macrophage gene expression.**

Each heatmap shows spearman correlation coefficients between averaged gene expression of each cluster in the placental dataset and that of annotated cell types from three single cell RNA-seq datasets.

**A.** 6-11 week placental signatures from Suryawanshi et al.[29]

**B.** 6–14 week placental signatures from Vento-tormo et al.[30]

**C.** Full term (≥37 week) placental signatures from COVID-19 negative control placentas, Lu-Culligan et al.[42] Black boxes indicate the best matched reference cell cluster for each of the HBC cell clusters identified in this work.
